# Large propulsion demands increase locomotor learning at the expense of step length symmetry

**DOI:** 10.1101/372425

**Authors:** Carly J. Sombric, Jonathan S. Calvert, Gelsy Torres-Oviedo

## Abstract

There is a clinical interest in increasing the extent of locomotor learning induced by split-belt treadmills that move each leg at different speeds. However, factors facilitating locomotor learning are poorly understood. We hypothesized that augmenting the braking forces, rather than propulsion forces, experienced at the feet would increase locomotor adaptation and learning. To test this, forces were modulated during split-belt walking with distinct slopes: inclined (larger propulsion than braking), declined (larger braking than propulsion), and flat (similar propulsion and braking). These groups were compared using step length asymmetry, which is a clinically relevant measure robustly adapted on split-belt treadmills. Unexpectedly, the group with larger propulsion demands (i.e., the incline group) adapted faster and more, and had larger after-effects when the split-belt perturbation was removed. We also found that subjects who propelled more during baseline and experienced larger disruptions of propulsion forces in early adaptation exhibited greater after-effects, which further highlights the catalytic role of propulsion on locomotor learning. The relevance of mechanical demands on shaping our movements was also indicated by the steady state split-belt behavior, during which each group recovered their baseline leg orientation to meet leg-specific force demands at the expense of step length symmetry. Notably, the flat group was nearly symmetric, whereas the incline and decline group overshot and undershot symmetry, respectively. Taken together, our results indicate that forces propelling the body facilitate gait adaptation during split-belt walking. Therefore, interventions that increase propulsion demands may be a viable strategy for augmenting locomotor learning in individuals with hemiparesis.

**Key Points Summary:**

- Split-belt walking (i.e., legs moving at different speeds) can induce locomotor learning and even improve hemiparetic gait, but little is known about how to facilitate this process.
- We investigated the effect of braking and propulsion forces on locomotor learning by testing young unimpaired subjects on the split-belt condition at different slopes (i.e., flat, decline, and incline), which distinctively modified these forces.
- Propulsion forces facilitated locomotor learning indicated by 1) greater adaptation and after-effects following split-belt walking of the inclined group, which experienced larger propulsion demands and 2) a positive correlation between individual after-effects and subject-specific propulsion during regular walking and initial steps in the split condition.
- Interestingly, incline and decline groups self-selected asymmetric step lengths at steady state in the split condition, challenging the prominent view that step length asymmetry is a biomarker for inefficient gait.
- Our results suggest that interventions augmenting propulsion demands could correct hemiparetic gait more effectively.

## Introduction

There is an interest in increasing the extent of locomotor adaptation and learning induced by split-belt walking because it can correct the step length asymmetry of individuals with hemiparesis, such as stroke survivors. Post-stroke rehabilitation targets step length asymmetry because it impairs mobility by augmenting the effort to walk (Waters & Mulroy, 1999), reducing walking speed (Balasubramanian *et al.*, 2007), compromising balance (Lewek *et al.*, 2014), and if not corrected, step length asymmetry leads to other comorbidities such as musculoskeletal injuries (Jørgensen *et al.*, 2000) and joint pain (Patterson *et al.*, 2012). Promising studies have shown that walking with the legs moving at different speeds (i.e., split-belt walking) results in long-lasting reduction of step length asymmetry post-stroke when walking overground (Reisman *et al.*, 2009, 2013). However, split-belt walking is not always effective even after repeated exposure to the split-belt experience (Reisman *et al.*, 2013; Lewek *et al.*, 2017) and it is still unclear why some stroke survivors re-learn to walk symmetrically but others do not. Thus, it is fundamental to identify factors facilitating split-belt adaptation to augment them for increasing motor learning in all individuals.

The increased mechanical work (Selgrade *et al.*, 2017), step length asymmetry (Reisman *et al.*, 2005), and hence metabolic effort (Finley *et al.*, 2013), upon introducing the split-belt environment are thought to drive locomotor adaptation. Notably, these three factors are large during the initial steps of split-belt walking and are minimized as subjects learn to walk in the split-belt context (Finley *et al.*, 2013; Selgrade *et al.*, 2017). Thus, modulating anterior-posterior forces applied at the feet during split-belt walking could facilitate locomotor adaptation given their direct impact on mechanical work and step lengths in regular gait (Donelan *et al.*, 2002). In particular, we hypothesize that altering braking forces could modulate the adaptation of gait based on prior studies showing that braking forces are critical for gait stability (Beschorner *et al.*, 2016) as subjects are most likely to fall during the braking phase of gait (Redfern *et al.*, 2001) and that braking forces are tightly regulated during split-belt walking (Ogawa *et al.*, 2014). Notably, braking forces are suddenly disrupted when the split-belt perturbation is introduced and this perturbation is reduced as subjects adapt their gait. Thus, we proposed that braking forces could facilitate subject-specific gait adaptation and therefore, increasing these forces would augment the extent of locomotor learning.

To test this hypothesis, we assessed subjects’ gait before, during, and after split-belt walking at different inclinations, which naturally and distinctively modulated the braking and propulsion forces experienced at the feet (Lay *et al.*, 2006, 2007). We found that inclination increased locomotor adaptation and learning as hypothesized, but not through the proposed mechanism. While the braking and propulsion forces were adapted and stored, it was the propulsion forces that were responsible for augmenting locomotor adaptation and learning, as quantified by the larger changes in step length asymmetry during and after split-belt walking. Specifically, declined walking that accentuated braking did not lead to as much adaptation as inclined walking, which augmented propulsion. In addition, propulsion forces during baseline and early adaptation were highly predictive of the extent of locomotor learning at an individual level. Interestingly, each inclination group recovered their baseline leg orientation to meet force demands at the expense of step length symmetry, which was surprising given the consistent human tendency to self-select symmetric step lengths in the split-belt environment (Bruijn *et al.*, 2012; Yokoyama *et al.*, 2018). Taken together, our findings suggest that propulsion demands, rather than braking forces, facilitate locomotor learning induced by split-belt walking.

## Methods

### Participants and Ethical Approval

We investigated the effect of modulating anterior-posterior forces applied at the feet on gait adaptation during and after split-belt walking (i.e., after-effects) under distinct slopes (i.e., flat, decline, incline), which naturally altered braking and propulsion forces (Lay *et al.*, 2006, 2007). To this end, we evaluated the kinetic and kinematic adaptation and after-effects of twenty four young healthy subjects (12 men and 12 women, 24.5±4.9 years of age) randomly assigned to one of three groups experiencing the split-belt adaptation protocol in a flat, incline, or decline configuration (n=8 each). The University of Pittsburgh Institutional Review Board approved the experimental protocol and conformed to the standards set by *Declaration of Helsinki*, except for registration in a database.

### General Paradigm

All subjects experienced the same split-belt paradigm illustrated in Figure 1A. Only the inclination at which subjects walked was altered across groups to determine the role of anterior-posterior ground reaction forces on the adaptation and learning of kinetic and kinematic gait features. We specifically tested subjects at three inclinations: flat (0 degrees), incline (8.5 degrees), or decline (−8.5 degrees). Subjects from each group walked at the specified inclination throughout the experiment, including baseline and post-adaptation.

**Figure 1.**
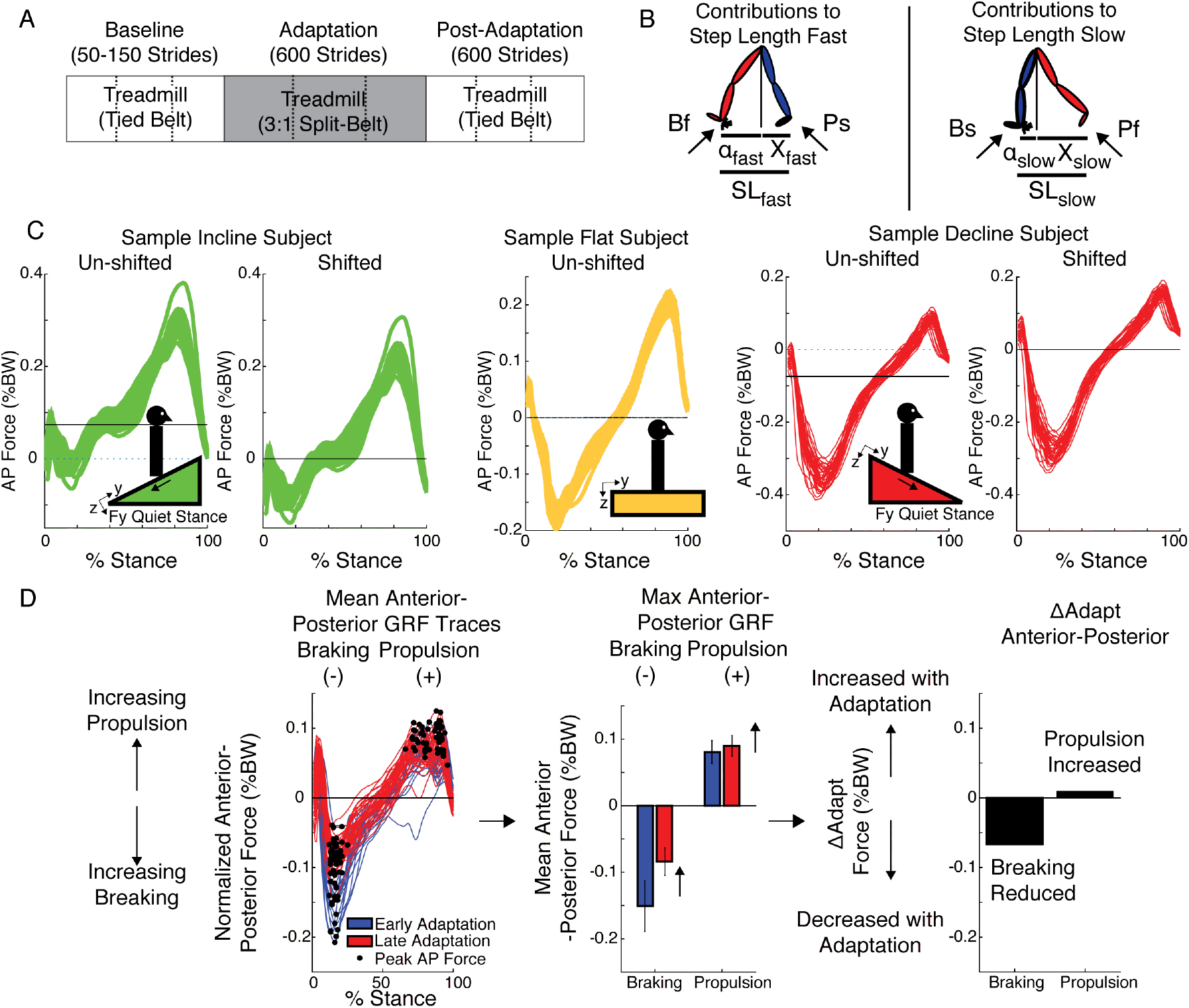
Experimental Paradigm and Kinetic Analysis. (A) Split-belt treadmill paradigm used for all sloped conditions to assess locomotor adaptation and learning of a new gait pattern. Subjects from each group walked either inclined (8.5°), flat, or declined (−8.5°) throughout the experiment, including baseline and post-adaptation. All groups experienced the same number of strides per epoch (baseline: between 50 and 150, adaptation: 600, post-adaptation: 600). Dashed lines indicate when the resting breaks occurred (B) The decomposition of step length into leading (α) and trailing (X) leg positions with respect to the body are illustrated. This decomposition was done because it is known that inclination affects these aspects of step length differently (Leroux, et al 2002; Dewolf et al 2017). Also note that when taking a step, the step length will depend on the position of the leading and trailing leg, which are generating a braking and propulsion force, respectively. (C) Anterior-posterior forces are plotted for sample subjects walking at the baseline epoch in each inclination condition. The filtered forces (un-shifted panels) clearly illustrate the effect of inclination on braking and propulsion forces: the incline group reduced braking and accentuated propulsion, the flat group experienced similar braking and propulsion, and the decline group accentuated braking and reduced propulsion. To facilitate the identification of peak braking (maximum negative value) and peak propulsion forces (maximum positive value) in all conditions, we removed the slope-specific projection of subjects’ weight on the AP direction (shifted panels). This procedure also allowed us to characterize the magnitude of kinetic outcome measures for each condition beyond those due to the slope-specific bias. (D) We used the peak braking and peak propulsion force for each step to compute outcome measures of interest, such as the ∆Adapt measure. This measure was computed to quantify increments or reductions in magnitude within the adaptation epoch of each specific parameter. Note that increases in magnitude were defined as positive changes, whereas decreases in magnitude were defined as negative changes.

#### Baseline

Subjects first experienced a baseline epoch to characterize their gait at the specific inclination at which they were going to walk throughout the study. During the baseline epoch, subjects walked with the belts moving at the same speeds (i.e., tied condition). Belts moved either at a slow (0.5m/s), fast (1.5 m/s), and medium (1 m/s) speeds for at least 50 strides. Strides were counted in real-time using raw kinetic data. A stride was defined as the period between two consecutive heel strikes (i.e., foot landing) of the same leg. Heel strikes were identified when the raw normal force under each foot reached a threshold of 30 Newtons.

#### Adaptation

the adaptation epoch was used to assess subjects’ ability to adapt to a new locomotor pattern in response to a split-belt perturbation. During this period, one leg moved three times faster than the other (0.5 m/s and 1.5 m/s) for 600 strides, except for one subject in the decline group who experienced the adaptation condition for 907 strides (due to technical difficulties in heel strike detection). This subject was not excluded from the analysis given that this participant’s behavior did not differ from that of other subjects and its inclusion did not alter the conclusions drawn from our results. The leg walking fast was the dominant leg, which was determined as the self-reported leg used to kick a ball.

#### Post-Adaptation

this epoch was used to assess the amount of locomotor learning when the split-belt condition was removed. Subjects experienced tied walking at the medium speed (1m/s) for 600 strides at their group-specific slope. Subjects where given resting breaks every 200 strides during the adaptation and post-adaptation periods to prevent fatigue.

### Data Collection

Kinematic and kinetic data were used to characterize subjects’ ability to adapt their gait during adaptation, and maintain the learned motor pattern during post-adaptation.

#### KinematicData

Kinematic data were collected with a passive motion analysis system at 100 Hz (Vicon Motion Systems, Oxford, UK). Subjects’ behavior was characterized with passive reflective markers placed symmetrically on the ankles (i.e., lateral malleolus) and hips (i.e., greater trochanter) and asymmetrically on shanks and thighs (to differentiate the legs). The origin of the kinematic data was rotated with the treadmill in the incline and decline conditions such that the z-axis (‘vertical’ in the flat condition) was always orthogonal to the surface of the treadmill (Figure 1C). Gaps in raw kinematic data were filled with a quintic spline interpolation (Woltring; Vicon Nexus Software, Oxford Uk).

#### Kinetic Data

Kinetic data was collected with an instrumented split-belt treadmill at 1,000 Hz (Bertec, Columbus, OH). Force plates were zeroed prior to each testing session so that each force plate’s weight did not affect the kinetic measurements. In addition, the reference frame was rotated at the inclination of each specific experiment such that the anterior-posterior forces were aligned with the surface on which the subject walked. Force data was low pass filtered at 20Hz.

### Data Analysis

#### Kinematic Data Analysis

Kinematic behavior was characterized with step length asymmetry, which exhibits robust adaptation in split-belt paradigms (e.g., Reisman *et al.*, 2005). It is calculated as the difference in step length between the two legs on consecutive steps. Step length is defined as the distance in millimeters between the ankle markers at heel strike. Therefore, equal step lengths result in zero step length asymmetry, whereas different step lengths result in non-zero step length asymmetry. A positive step length asymmetry indicates that the fast leg’s step length (which for this study is the dominant leg’s step length) is longer than the slow leg’s step length. Step length asymmetry was normalized by stride length (SL), which is the sum of two consecutive step lengths, resulting in a unitless parameter that is robust to inter-subject differences in step sizes.

Each step length was also decomposed into anterior and posterior components relative to the hip position (Figure 1B) as in previous work (Finley *et al.*, 2015). This was done because inclination is known to affect these measures (Leroux *et al.*, 2002; Dewolf *et al.*, 2017). The anterior component, ‘α’, was defined as the distance in millimeters between the leading leg’s ankle and the hip at heel strike; similarly, the posterior component, ‘X’, was defined as the distance in millimeters between the trailing leg’s ankle and the hip. The hip position, which is a proxy for the body’s position, was estimated as the mean instantaneous position across hip markers. α-distances were positive because the foot usually lands in front of the hips, whereas X-distances were negative because the trailing leg is behind the hips at foot landing. Note that the summation of the magnitudes of α and X equals the leading leg’s step length.

#### Kinetic Data Analysis

We characterized how ground reaction forces were modulated by inclination. The kinetic analysis was focused on forces in the anterior-posterior direction since these are modulated by inclination (e.g., Lay *et al.*, 2006) and they are adapted during split-belt walking (Ogawa *et al.*, 2014). The anterior-posterior ground reaction forces (AP forces) where normalized by each subject’s body weight to account for inter-subject differences in weight. The quiet stance forces for the incline and decline groups were subtracted in order to facilitate the identification of both positive (propulsion) and negative (braking) features of the AP forces at all slopes and to characterize the magnitude of these force for each condition beyond those due to the slope-specific bias. The removal of the quiet stance forces is illustrated for a representative subject at each inclination in Figure 1C. Note that at quiet stance (i.e., when subjects are just standing), there is an anterior-posterior ground reaction force due to the projection of subjects’ weight on the surface of the treadmill. This quiet stance force is zero for the flat group since AP forces are negligible during quiet stance on a level surface. However, this quiet stance force was substantial for the inclined and declined group. Thus, we calculated the quiet stance force based on the inclination of the treadmill and subtracted it from AP forces during gait (Eq. 1).

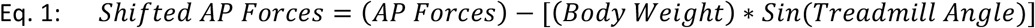

The shifted AP forces were further decomposed into braking and propulsion forces for each stride. The braking force was defined as the component of the shifted AP force with negative values. The braking forces were computed independently for the slow (BS) and fast legs (BF). On the other hand, the propulsion force was defined as the component of the shifted AP force with positive values that followed the braking phase (Figure 1D). The propulsion forces were also computed independently for the slow (PS) and fast legs (PF). We focused our analysis on the peak braking and propulsion forces for each leg and for each stride (Figure 1D, first panel) to be consistent with prior split-belt studies (Mawase *et al.*, 2013; Ogawa *et al.*, 2014) and those reporting kinetic differences between inclinations (Lay *et al.*, 2006; Item-glatthorn *et al.*, 2016).

### Kinetic and Kinematic Outcome Measures

Outcome measures were used to characterize the adaptation and after-effects of kinematic and kinetic gait features in response to a split-belt perturbation. A comprehensive list of outcome measures is provided in Table 1. Medium baseline behavior was used as a reference in all outcome measures computed with kinematic parameters (e.g., step length asymmetry and step lengths), whereas speed-specific baselines were used for kinetic parameters (e.g., braking and propulsion forces). In other words, fast baseline was used as a reference for the leg walking fast during adaptation and the slow baseline was used as a reference for the leg walking slow during adaptation, whereas medium baseline was used as a reference for both legs when they walked at the same medium speed in post-adaptation. This methodology is consistent with prior split-belt studies indicating that kinetic parameters plateau near values similar to those of the speed-specific baseline (Ogawa *et al.*, 2014). Outcome measures of interest were Early Adaptation, Late Adaptation, After-Effects, ∆Adapt, and ΔPost. **Early Adaptation** (EarlyA) was defined as the difference between the averaged behavior of the first 5 strides of the split-belt condition (strides 1-5) and the speed-specific baseline values as indicated earlier. This outcome measure characterized the extent to which subjects were perturbed by the split-belt condition. Note that we did not exclude any strides of adaptation because all subjects experienced a short split condition before the adaptation period to minimize startle effects. **Late Adaptation** (LateA) was defined as the average of the last 40 strides of adaptation for all parameters. This outcome measure indicated the steady state behavior reached at the end of the adaptation period. **After-Effects** were defined as the average of the first 5 strides of Post-Adaptation relative to medium baseline such that increments and reductions in magnitude of a specific parameter with respect to medium baseline were marked as positive or negative, respectively. We also characterized the behavioral changes within the adaptation and post-adaptation with indices ∆Adapt and ∆Post, respectively. ∆**Adapt** and ∆**Post** were computed as the difference between Late and Early Adaptation for ∆Adapt and late and early post-adaptation for ∆Post (i.e., average of the last 40 strides or 5 strides for late and early, respectively). This was done such that an increase in the magnitude of a parameter during either adaptation or post-adaptation resulted in positive values and a reduction of the parameter was marked as negative values. For example, we illustrate ΔAdapt for braking and propulsion forces in Figure 1D. Note that the braking force decreased during the adaptation period (negative value), whereas the propulsion force increased during the same period (positive value). Lastly, the effect of slope on the rate of adaptation was determined by fitting the averaged step length asymmetry for each inclination group with a single exponential (y=a*exp((−1/τ)*x)+c) using a non-linear least squares method. The 95% confidence intervals of the Tau coefficient were used to compare the adaptation rates across groups.

**Table 1:** Outcome Measures

| Outcome Measure | Meaning |
| --- | --- |
| Early Adaptation (EarlyA) | Quantifies how perturbing split-belt walking is for a given parameter. This parameter is reported in reference to medium baseline for symmetry parameters, and in reference to speed specific baseline for leg-specific parameters. |
| Late Adaptation (LateA) | Quantifies the amount that the split-belt perturbation was counteracted for step length asymmetry only. This parameter is reported in reference to medium baseline for symmetry parameters. |
| After-Effects | Quantifies how much motor learning, or how much of the split-belt walking pattern subjects use following adaptation relative to medium baseline behavior. Positive values indicate that the post-adaptation is larger in magnitude than if was during baseline, except for individual step lengths where positive values indicate that the step length is increased in the Post-Adaptation Epoch. |
| $\Delta\text{Adapt}$ & $\Delta\text{Post}$ | Quantifies the change in a parameter during adaptation and post- |
| | adaption, respectively. Therefore, $\Delta\text{Adapt}$ and $\Delta\text{Post}$ enabled us to determine if changes during adaptation led to comparable changes during post-adaptation. Positive means there was an increase in magnitude of a parameter within an epoch and vice versa. |
| Rate of Adaptation | Quantified the group mean rate of adaptation of step length asymmetry for during adaptation assuming a single exponential. |
| Baseline Propulsion Magnitude | Quantifies subject's baseline propulsion force magnitude characteristic for each leg during medium baseline, which is the average walking speed during adaptation. |
| Baseline Propulsion Asymmetry | Quantifies subject's propulsion force laterality in the most demanding baseline condition (i.e., fast baseline). |

### Post-Hoc Data Analysis

Following our planned analysis, it was clear that propulsion forces were more important for locomotor adaptation and learning processes than anticipated. Thus, features characterizing each subject’s propulsion forces were used as predictors in a statistical model to determine after-effects of step length asymmetry at an individual level. In this propulsion-based model, we hypothesized that perturbation of the propulsion forces per leg (i.e., propulsion during early adaptation) would be predictive of step length asymmetry after-effects. We also considered that each individual’s magnitude of baseline propulsion could influence after-effects because we reasoned that those with larger propelling tendencies maybe more prone to correcting reductions to their propulsion forces. Additionally, we included subject-specific propulsion asymmetries during baseline in our model because we thought that those who are naturally more asymmetric could be less resistant to update their movements for achieving the large propulsion asymmetry required for split-belt walking. In sum, a multiple regression analysis was performed with step length asymmetry after-effects as a dependent variable and 6 regressors: the propulsion force perturbation of each leg, the baseline capacity to propel of each leg, the asymmetry of the baseline propulsion forces and a categorical factor indicating the inclination condition. Every subject’s Baseline Propulsion Magnitude for each leg was computed during medium baseline, which is the average belt speed during adaptation and post-adaptation epochs. On the other hand, each subject’s Baseline Propulsion Asymmetry was computed as the difference in average behavior between the dominant and non-dominant legs during the most challenging baseline condition (fast baseline), which increased this measure of kinetic asymmetry facilitating its assessment. We specifically computed the mean difference between the dominant minus the non-dominant propulsion during fast baseline. Therefore, zero indicated that both legs generate equal propulsion forces, positive numbers indicated that the dominant leg propelled more than the non-dominant leg and vice versa for negative numbers. We also perform two additional analyses to evaluate the predictive power of the propulsion-based model on after-effects of step length asymmetry at an individual level. First, we applied the same procedure described above using step length, rather than propulsion force, to determine if any information about the behavior of individual legs was predictive of the inter-subject variability of after-effects in step length asymmetry. Second we used a mean-based model that only included the categorical factor as a regressor to test if the propulsion-based model accounted for more inter-subject variability on step length asymmetry after-effects than the group mean behavior. The explanatory power between models was contrasted with Bayes Factor Analysis (Kass *et al.*, 1995) using the Bayesian Information Criteria approximation (Wagenmakers, 2007). Lastly, several multiple linear regressions between kinetic and kinematic variables were performed within and across epochs to investigate the association between propulsion forces and after-effects in step length asymmetry. Of note, the regression of propulsion forces across epochs (i.e., between ∆Adapt and after-effects) was done with contralateral legs to be consistent with the post-hoc observation that adaptation of step length on one side led to after-effects on the other side (see Figure 3).

### Statistical Analysis

One-way ANOVAs were used to test the effects of inclination condition on kinetic and kinematic outcome measures (e.g., ∆Adapt, After-Effects, etc.). Outcome measures with significant ANOVAs were further analyzed with Tukey post-hoc testing. We compared the time constants estimated from single exponential fits of group averages of step length asymmetry using the 95% confidence intervals for each rate coefficient. Time constants were determined to be significantly different when the confidence intervals were not overlapping. We additionally wished to know if each group’s step length asymmetry steady state was different from zero, therefore we performed a two-sided t-tests on each group’s late adaptation values. A three-way ANOVA was also performed to determine the effect of leg and inclination condition on the changes of step length that occurred during adaptation and post-adaptation. As such, the dependent variables were changes of step length that occurred during these epochs (i.e., ∆Adapt and ∆Post); the independent variables were slope (i.e., incline, flat, decline), leg (i.e., fast or slow leg), epoch (i.e., adaptation or post-adaptation), and the interactions between these independent variables. Lastly, two multiple regressions were performed for step length asymmetry to assess the association between 1) the perturbation (EarlyA) vs. adaptation (∆Adapt) of this parameter and 2) its adaptation vs. its after-effects. In these regressions group was used as a categorical factor only if it was found to be independent of the continuous variable (i.e., EarlyA for regression 1 and ∆Adapt for regression 2) as indicated by a non-significant Pearson Coefficient. Should a strong correlation between the categorical and continuous regressor was found, a linear regression was performed and the confounding influence of group was noted. For visualization purposes only, we displayed the results of a linear regression when the continuous variable of the multiple regression analysis was a significant factor. A significance level of α=0.05 was used for all statistical tests. MATLAB was used for all statistical analysis (The MathWorks, Inc., Natick, Massachusetts, United States).

## RESULTS

### Inclination regulated the adaptation and after-effects of step length asymmetry

Step length symmetry was not recovered in the sloped split-belt conditions. All groups were perturbed by split-belt walking and subsequently adapted (Figure 2A). However, each group reached a different step length asymmetry by Late Adaptation (p<0.001) such that the flat group plateaued near symmetric step lengths (SLA_LateA=0, p=0.08), whereas the decline group undershot step length symmetry (SLA_LateA<0, p<0.001) and the incline group overshot step length symmetry (SLA_LateA >0, p<0.001) (Figure 2B). Interestingly, while both sloped groups were perturbed more than the flat group (SLA_EarlyA: group main effect p=0.003; incline vs. flat p=0.036; decline vs. flat p=0.002), only the incline group adapted more than the other two groups (SLA_∆Adapt: group main effects p<0.001; incline vs. flat p<0.001; incline vs. decline p=0.002) and faster (Figure 2A; inlayed plot). Importantly, groups were not only distinct in their adaptation, but also in their after-effects (p=0.002) such that the incline group had greater after-effect than the decline (p=0.03) and flat groups (p=0.002). Our regression analyses on individual subjects revealed that larger perturbations led to more adaptation (pmdl<0.001, R^2^=89, SLA_EarlyA: p<0.001) and greater adaptation resulted in more after-effects (R^2^=46; p<0.001). However, these associations were likely driven by group differences since group was either a significant categorical predictor of adaptation (Group: p<0.001) or group was strongly correlated to the continuous predictor of after-effects (p=0.004). In other words, the larger perturbation and adaptation of the inclined condition presumably resulted in greater after-effects in this group compared to others. Therefore, contrary to our hypothesis, the slope condition augmenting propulsion (i.e., incline walking) rather than braking led to more and faster adaptation and greater after-effects. It was also unexpectedly found that the sloped groups plateaued at values that were distinct from their baseline step length symmetry, begging the questions of whether inclination had a different effect across individual step lengths (addressed in Figure 3) and why this would happen (addressed in Figure 4).

**Figure 2.**
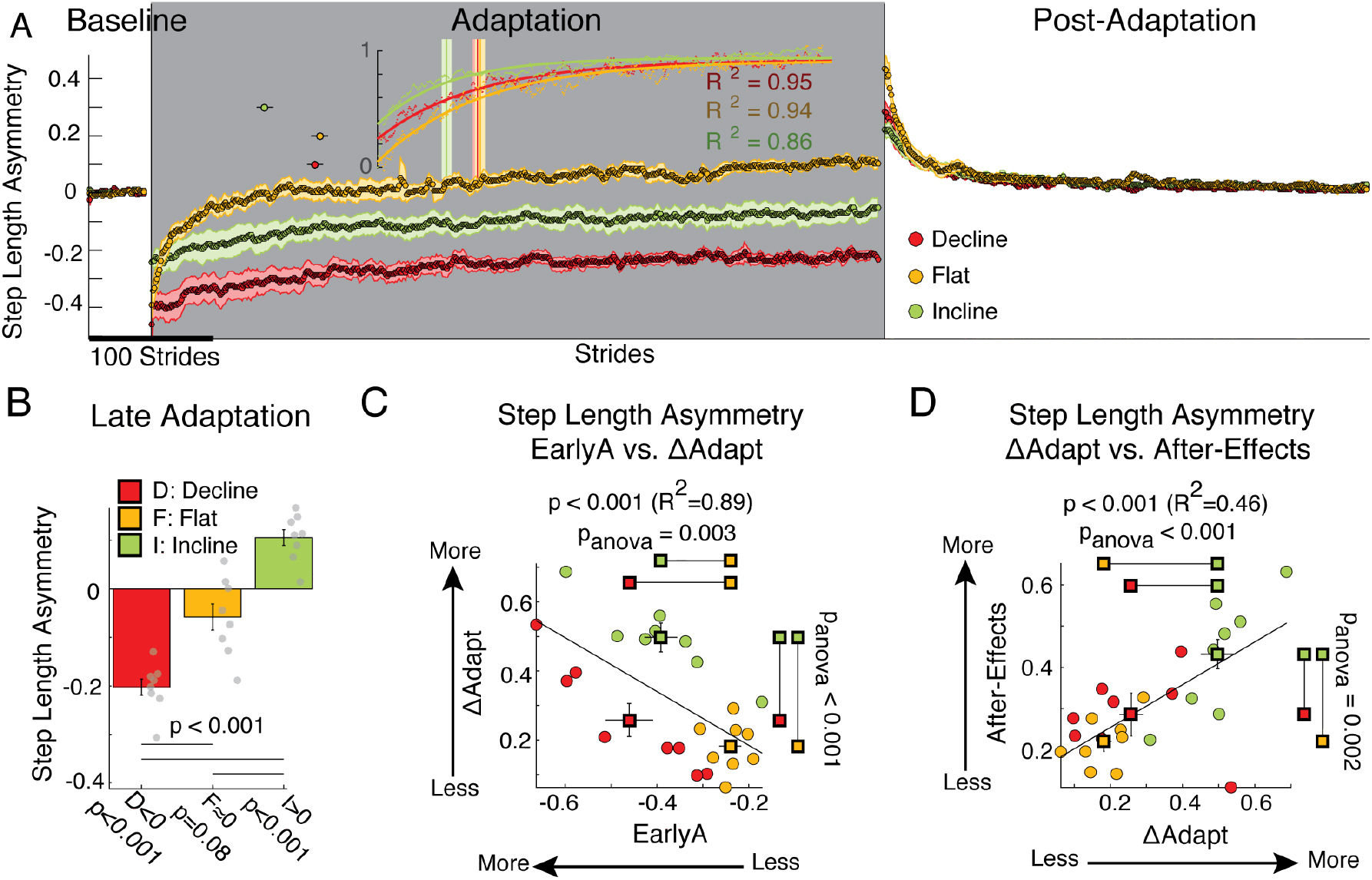
Step length Asymmetry Adaptation & Learning. (A) Stride-by-stride time course of step length asymmetry during medium baseline, adaptation, and post-adaptation are shown. Each data point represents the average of 5 consecutive strides and shaded regions indicate the standard error for each group. Note that the incline group adapted more quickly than the other sloped conditions as indicated by the time constants found by fitting the group data with a single exponential (R^2^>=0.86). The time constants are plotted with 95% confidence intervals during the adaptation phase and the inlayed axis shows the fits of the adaptation timecourses scaled between 0 (Early Adaptation) and 1 (Late Adaptation) for each group. (B) Bar plots indicate the averaged step length asymmetry for each group during Late Adaptation ± standard errors and horizontal lines indicate statistical differences between groups according to post-hoc testing. Note that each group plateaued at different step length asymmetry values that were statistically different from symmetry (zero value) as indicated by individual t-tests reported below the x-axis. Individual subject behavior is indicated with grey dots for each group. (C) Results from multiple regression analysis between ∆Adapt and two uncorrelated regressors: slope condition and step length asymmetry during early adaptation. The regression model was significant pmdl<0.001, R^2^=0.89 and both regressors were significant predictors. In panel C and D, values for individual subjects are illustrated with colored dots and mean values for the group are illustrated with colored squares. Lines on the squares represent standard error bars. p_anova_ parallel to either the x-axis or y-axis indicates the effect of inclination condition on the outcome measure plotted on the respective axis. For example, panel C shows a significant group effect on step length asymmetry during Early Adaptation (p_anova_=0.003) and delta (p_anova_<0.001). In addition, significant post-hoc differences between groups are illustrated with lines connecting colored squares. For example, panel C also illustrates that although the decline (red square) and incline groups (green square) were more perturbed by split-belt walking than the flat group (yellows square), only the incline group (green square) adapted more than the other two groups. (D) Linear regression between ΔAdapt and After-Effects for step length asymmetry was significant (p<0.001, R^2^=0.46). This indicated that the more step length asymmetry was adapted (large ΔAdapt values), the greater the after-effects. However, this association was strongly influenced by group differences. Specifically, the incline group who adapted the most, had the largest step length asymmetry after-effects.

**Figure 3.**
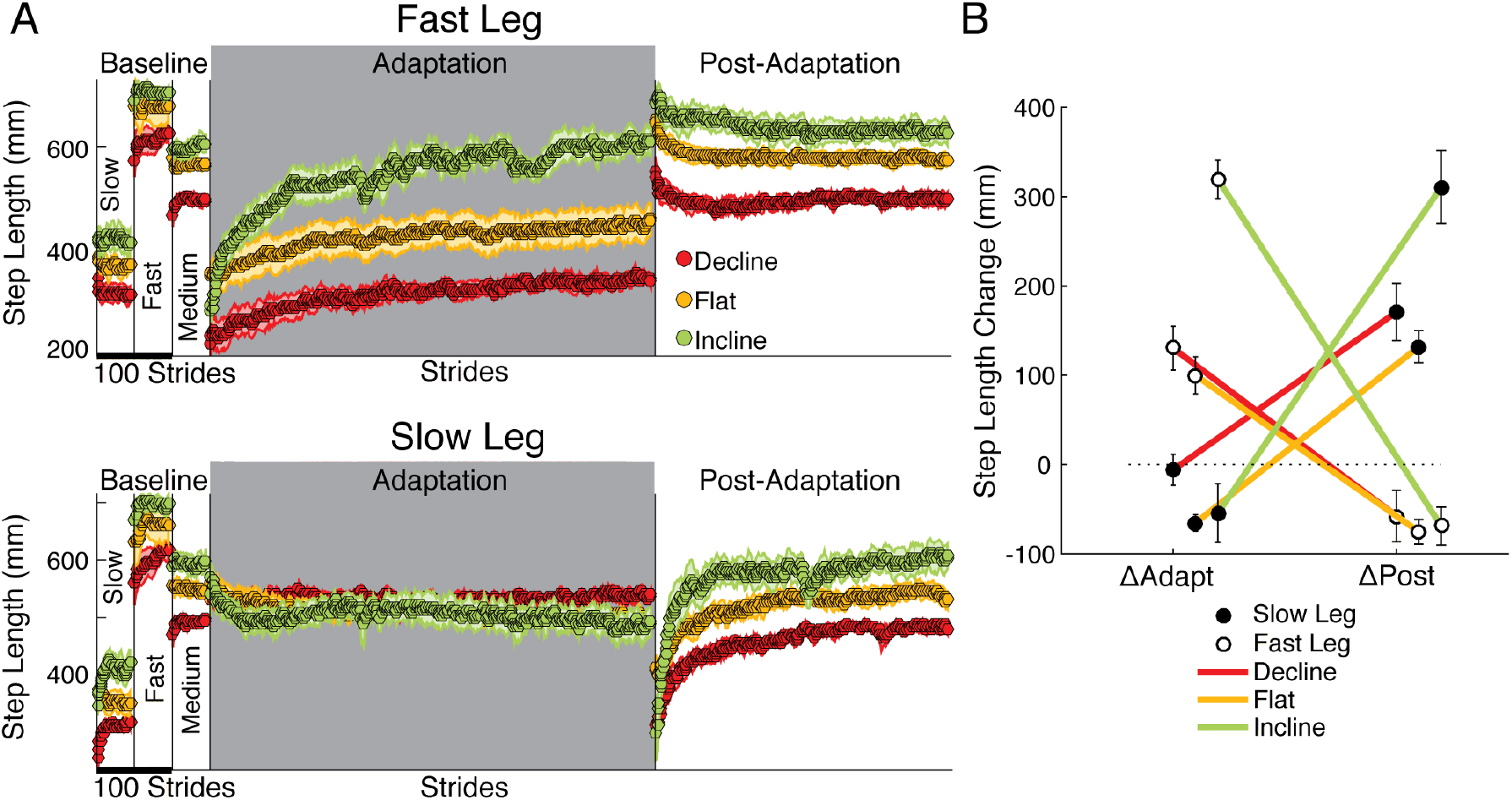
Step length Adaptation & Learning. (A) Stride-by-stride time courses of both step lengths are shown during slow baseline, fast baseline, medium baseline, adaptation, and post-adaptation. The medium baseline bias was not removed to appreciate the distinct step length values across sloped conditions. Each data point represents the average of 5 consecutive strides and shaded regions indicate the standard error for each group. Step lengths were different across groups in all baseline speeds. In addition, the fast step length of the inclined group exhibited the largest change during the adaptation period (largest ∆Adapt across groups). In contrast, the slow step length was similarly adapted across groups and approached the same value at steady state in all inclination conditions (p=0.17 in OneWay ANOVA of biased values during Late Adapt). On the other hand, the slow step length had large and slope-mediated after-effects, whereas the fast step length had modest and similar after-effects across sloped conditions. (B) The effect of slope on each leg’s change during adaptation (∆Adapt) and post-adaptation (∆Post) is illustrated. Note the contralateral relation between adaptation and post-adaptation in all sloped conditions: the fast step length adapted more than the slow one (large ∆Adapt for fast leg), whereas the slow step length had most of the after-effects. This contralateral relation is exaggerated by incline walking.

**Figure 4.**
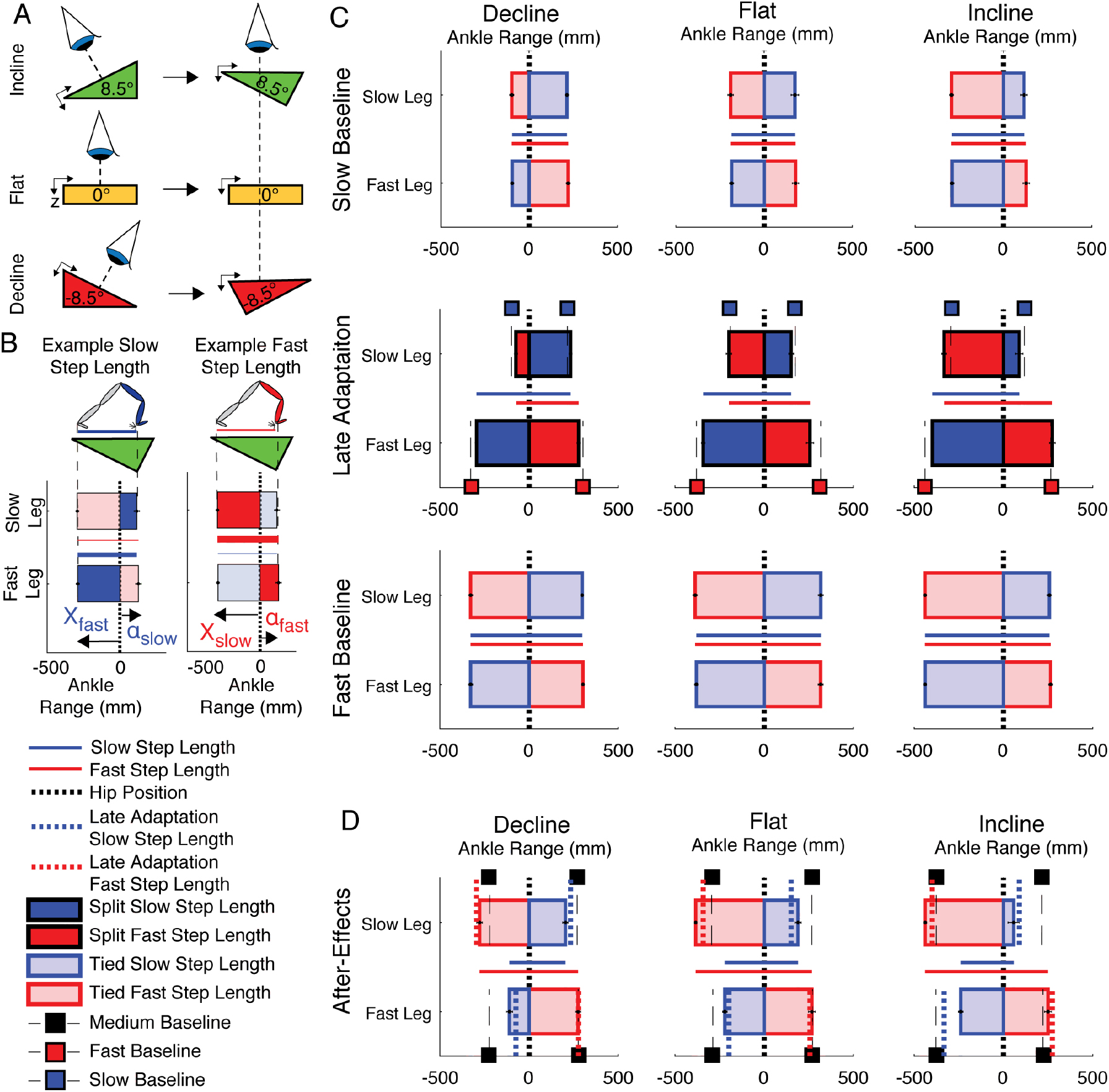
Leg orientation Adaptation & After-Effects. (A) Schematic indicating the view projected in panels B through D. (B) Visualization approach used in panels C and D. Vertical zero lines represent the perpendicular projection of the hips onto the treadmill. The horizontal bars represent the ankle positions with respect to the hips at ipsilateral and contralateral heel strikes ± standard errors. Each bar’s maximum positive value indicates the ankle position at ipsilateral heel strike (α-position), whereas the maximum negative value indicates the ankle position at contralateral heel strike (X-position). The horizontal lines plotted between the two horizontal bars represent the step lengths for when the fast leg is leading (in blue) or for when the slow leg is leading (in red). (C) Leg orientations are illustrated for both legs during baseline walking slow (top row), baseline walking fast (bottom row), and late adaptation (middle row). Vertical thin dashed lines are plotted to indicate the α-position and X-position during slow baseline walking (dashed lies with red squares) or fast baseline walking (dashed lines with blue squares). We observed that the α-position and X-position during late adaptation in all sloped conditions (middle row) were similar to the baseline positions for each specific speed and inclination, as a result, step lengths were asymmetric in the sloped conditions. (D) Leg orientations are illustrated during early post-adaptation (after-effects) at each sloped condition. We also plotted the α-position and X-position during baseline walking at medium speed (dashed lines with black squares) and during late adaptation (thick dotted lines). Note the ipsilateral similarity between the leading leg’s position (α-position) when taking a step during late adaptation and post-adaptation. In contrast, there was a contralateral similarity between the trailing leg’s positions during late adaptation and post-adaptation. For example, X-position during post-adaptation of the leg that walked slow (solid red line) was similar to the X-position of the fast leg during late adaptation (dotted red line).

### Inclination accentuates the adaptation of step lengths on the fast leg and subsequent after-effects on the slow leg

While incline walking augmented the adaptation and after-effects of step length asymmetry, it had a surprising unilateral effect on individual step lengths: it predominantly increased the adaptation of one leg and the after-effects of the other one. Figure 3 illustrates the time courses for each step length in every group. While both legs adapted and had after-effects in all groups, the adaptation of the fast leg was greater than that of the slow one, whereas the opposite was observed during post-adaptation. In other words, the leg that was modified the most switched between the fast and slow legs depending on the epoch, which was substantiated by the significant interaction between leg and epoch on the changes of step length (Figure 3B: leg#epoch p<0.001). In addition, the sloped conditions augmented this switching between legs, as indicated by the significant interaction between leg, epoch, and slope (p<0.001). Once more, the incline group drove this effect by exhibiting the largest changes on the fast side during adaptation (∆Adapt: incline vs. flat p<0.001; ∆Adapt: incline vs. decline p<0.001) and on the slow side during post-adaptation (∆Post: incline vs. flat p=0.003; ∆Post: incline vs. decline p=0.02). The impact of inclination on the changes of step length was further substantiated by the significant effect of slope (p<0.001) in contrast with the non-significant effect of leg (p=0.13) or epoch (p=0.89). In sum, inclined walking exaggerated the split-belt phenomenon of predominantly adapting the step length of the fast leg and observing after-effects on the other leg during post-adaptation.

### Slope- and Speed-Specific walking demands determine the distinct step length asymmetry across inclination conditions

Leg orientation mediated by inclination and walking speed underlay the distinct step length asymmetries across sloped conditions. More specifically, subjects oriented their legs to prioritize slope-specific demands on AP forces to walk at the speed imposed under each leg at the expense of step length symmetry during adaptation. This is shown in Figure 4 illustrating the top view of ankle positions relative to the body (Figure 4A) and the step lengths to which these positions contribute (Figure 4B) at baseline walking (slow and fast speeds) and late adaptation under each sloped condition (Figure 4C). Note that in slow baseline walking (Fig. 4C 1^st^ row) subjects shifted their ankle position either forward (i.e., |α|>|X|) or backward (i.e., |X|>|α|) relative to their body in the decline and incline condition, respectively (Fig 4C decline: 1^st^ row, 1^st^ column; incline: 1^st^ row, 3^rd^ column). This was done to facilitate the generation of either braking or propulsion forces counteracting the increased acceleration of center of mass in the decline group and deceleration of center of mass in the incline group due to gravity. Importantly, these braking and propulsion demands were additionally regulated by walking speed, as shown by the distinct leg orientations between slow (Fig. 4C 1^st^ row) and fast baseline walking (Fig. 4C 3^rd^ row). These inclination- and speed-mediated demands persisted during split-belt walking. Consistently, subjects oriented their legs in late adaptation at all sloped conditions according to the speed at which each leg walked. In other words, the slow and fast legs’ orientation in late adaptation was similar to the one at slow and fast baseline walking, respectively (Fig 4C, 2^nd^ row: fast and slow leg orientation in the speed-specific baseline is indicated by red and blue dashed lines, respectively). These leg orientations consequently led to step lengths that were only the same at the end of adaptation in the flat group, but not in the two sloped conditions (Fig 4C, 2^nd^ row: fast and slow step lengths are shown by red and blue solid lines, respectively). Lastly, leg orientation during post-adaptation indicated that the forward (α) positions of the legs were roughly the same before and after removal of the split perturbation, which deferred from α-positions at medium baseline walking (Fig 4C. 4^th^ row, black dashed lines). Conversely, the trailing positions (X) were swapped between legs: X-position of the slow leg during post-adaptation was similar to the X-position of the fast leg during late adaptation (dotted red lines) and vice versa for the other leg. This switching contributed to the reduced step lengths of the slow leg during post-adaptation (Fig 4C. 4^th^ row blue solid lines) also presented in Figure 3. Thus, leg orientation did not immediately return to baseline behavior during early post-adaptation, indicating that leg orientation is not purely reflecting the state of the environment but it is adjusted through an active adaptive process. In sum, asymmetric step lengths in sloped split-belt walking resulted from the adaptation of leg orientation to generate the forces for walking at the specific speed and inclination imposed on each group, which will be further discussed in the following sections.

### Braking and propulsion forces were predominantly changed in the inclined and declined groups, respectively

We assessed the adaptation of forces under the distinct inclination conditions before investigating the relation between kinetic and kinematic measures during and after split-belt walking. We found that braking and propulsion forces were predominantly modified during adaptation and post-adaptation in the sloped condition that naturally prioritized them. In other words, braking was mostly modulated in the decline group, whereas propulsion was primarily modulated in the incline one. This preferential regulation of braking and propulsion forces was indicated by the significant group effect on early adaptation for braking forces on both legs Bs_EarlyA: p =0.003; Bf_EarlyA: p<0.001) and propulsion forces of the slow leg (Ps_EarlyA: p =0.038) but not the fast one (Pf_EarlyA: p_anova=0.15). More specifically, the braking and propulsion forces were respectively more perturbed in the decline and incline groups compared to the flat group (Bs_EarlyA: decline vs. flat p=0.015; Bf_EarlyA: decline vs. flat p<0.001; Ps_EarlyA: incline vs. flat p<0.001). Despite the group differences in early adaptation, only the forces contributing to the step lengths of the fast leg (i.e., fast breaking and slow propulsion) were adapted differently across groups (Bf_∆Adapt: p<0.001; Ps_∆Adapt: p <0.001), whereas the other forces were adapted similarly across sloped conditions (Bs_∆Adapt: p=0.10; Pf_∆Adapt: p=0.35). Once again braking was preferentially adapted in the decline group compared to flat walking (p=0.0011) and propulsion was preferentially adapted in the incline group compared to flat walking (p=0.0041). Group analyses on post-adaptation indicated that sloped condition had a significant impact on the after-effects of slow braking (p<0.001), slow propulsion (p=0.036), and fast propulsion (p<0.001) and a trending effect on fast braking (p=0.076). However, only the forces contributing to after-effects of the slow step length (i.e., slow braking and fast propulsion) exhibited greater after-effects than those regularly observed after split-belt walking in a level surface (Bs_∆Post: decline vs. flat p<0.001; Pf_∆Post: incline vs. flat p=0.006 in contrast with Ps_∆Post: incline vs. flat p=0.92). In sum, inclination conditions predominantly modified the perturbation, adaptation, and after-effects of braking and propulsion forces in a preferential manner: inclined walking mostly modified the adaptation of propulsion, whereas decline walking mostly altered the adaptation of braking.

### A propulsion-based model predicted subject-specific after-effects in step length asymmetry

While braking and propulsion forces were both altered with inclination during and after split-belt walking, the adaptation of movements only increased in the incline group who experienced greater propulsion demands. This suggested that propulsion forces, rather than braking ones, played an important role in the adaptation and after-effects of walking movements. Consistently, we found that a propulsion-based model could predict individual after-effects in step length asymmetry (F(4, 20)=17.5, R^2^=0.73, p<0.001) (Figure 6A). In particular, we observed a significant association between after-effects in step length asymmetry and the perturbation of the slow leg’s propulsion (t=-2.42, p=0.028), the slow leg’s baseline propulsion (t=2.38, p=0.030), and baseline’s propulsion asymmetry (t=2.88, p=0.011). These significant coefficients in the multiple regression equation indicated that subjects exhibiting greater after-effects in step length asymmetry were 1) those who were more perturbed during split-belt walking, 2) those who naturally had higher propulsion tendencies during baseline walking, and 3) those who naturally had larger propulsion asymmetries (i.e., larger propulsion with their dominant than non-dominant leg) during baseline walking. Interestingly, these associations were exclusive to the non-dominant (i.e., slow) leg given that propulsion features of the dominant (i.e., fast) leg during baseline (t=-0.47, p=0.64) or early adaptation (t=1.20, p=0.25) were not significant predictors of after-effects in step length asymmetry. Importantly, inclination (categorical factor) was also not a significant predictor (t<0.28, p>0.78) in the multiple regression analysis, indicating that the propulsion-based model was not simply reflecting the differences in after-effects across groups (reported in Figure 2). The non-significant predictive power of the sloped condition was reiterated by the Bayes Factor of 925 indicating that the propulsion-based model had “very strong” predictive power (Kass Raftery, 2007) compared to a model that only included the mean after-effect values for each group. This larger predictive power of the propulsion-based model was maintained, but to a lesser extent when using more strides to characterize after-effects (1:10 strides, BF=97.3) as in prior work (Malone *et al.*, 2012). Lastly, the propulsion-based model was still a better predictor of subject-specific step length asymmetries than a multiple regression model based on individual step lengths (F(2, 22)=6.04, R^2^=0.21, p=0.02; Bayes Factor of 11.9*10^3), in which only the disturbance of the slow step length was a significant factor (SLs_EarlyA: t=-2.26 p=0.038; SLf_EarlyA: t=-2.05, p=0.057; SLs_base: t=-0.20, p=0.84; SLf_base: t=-0.47, p=0.65; SLasym: t=-0.06, p=0.95). This indicated that subject-specific propulsion forces during baseline and adaptation provided more information about inter-subject variability of after-effects than step lengths of individual legs, which is remarkable given that step lengths were used to directly compute step length asymmetry. Therefore, each participant’s tendencies to regulate propulsion forces during baseline and adaptation epochs were a strong indicator of individual after-effects in step length asymmetry, which represent subject-specific learning during split-belt walking.

**Figure 5.**
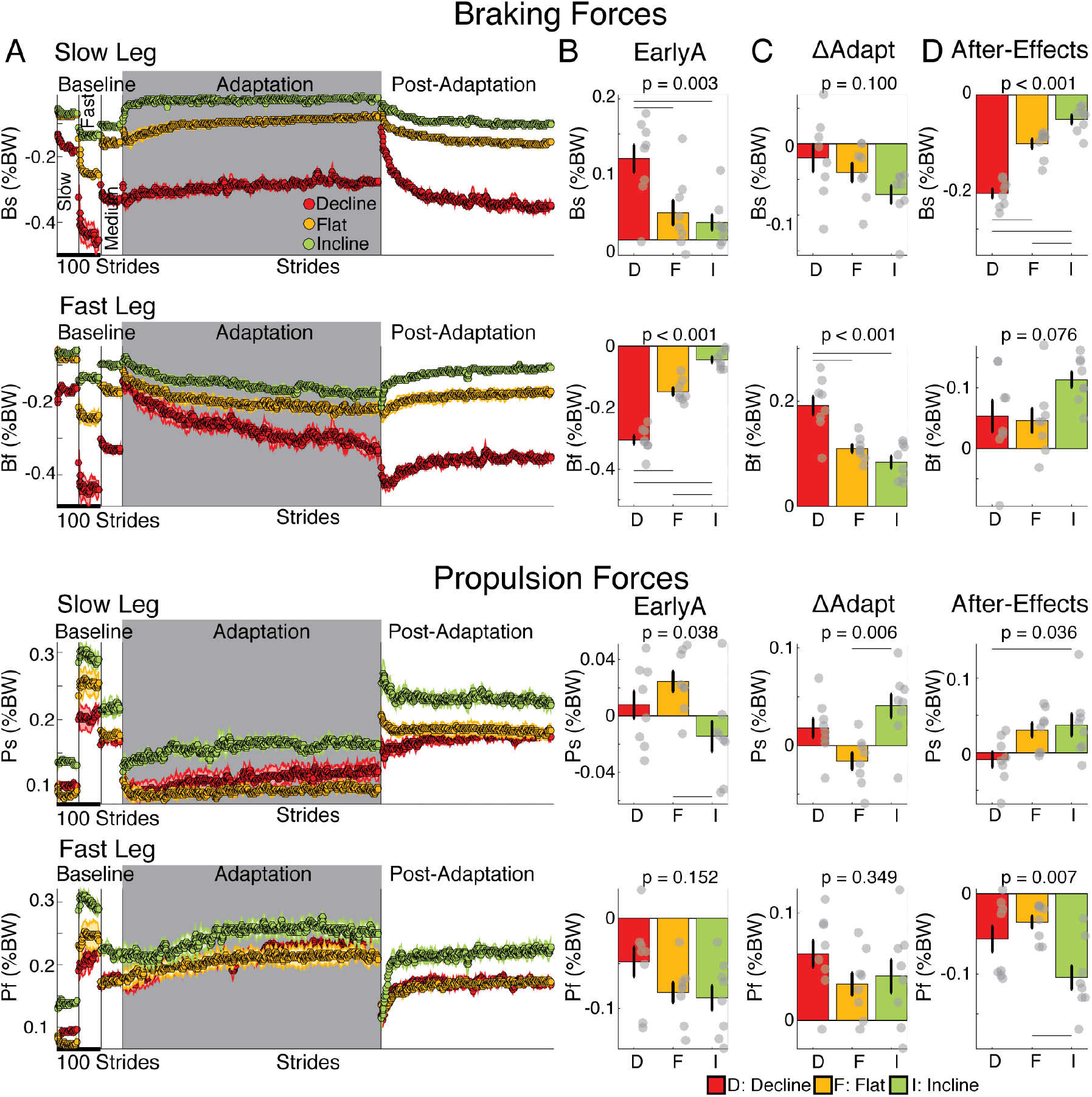
Adaptation and after-effects of braking and propulsion forces. (A) Stride-by-stride time courses of braking and propulsion forces by each leg are shown during slow, fast, and medium baseline, adaptation, and post-adaptation. Each data point represents the average of 5 consecutive strides and shaded regions indicate the standard error for each group. At the split-to-tied transition, braking forces changed smoothly, particularly for the flat (yellow) and incline (green) groups, but propulsion forces changed discontinuously in all groups. (B) **Early Adaptation:** The height of the bars indicates group averages for braking and propulsion forces for each leg during Early Adaptation forces ± standard errors. We observed that the braking and propulsion forces were more perturbed in the decline and incline groups, respectively, compared to the flat group. In panels B through D, positive and negative values respectively indicate increments or reduction in force relative to speed-specific baselines. Thin horizontal lines between groups illustrate significant differences (p<0.05) based on post-hoc analysis and gray dots indicate values for individual subjects. (C) ∆**Adapt:** The height of the bars indicates group averages for the adaptation of braking and propulsion forces for each leg during split-belt walking ± standard errors. We observed that the adaptation of braking was significantly greater in the decline than the flat group, whereas the adaptation of propulsion was significantly larger in the incline than the flat group. (D) **After-Effects:** The height of the bars indicates group averages After-Effects during early post-adaptation relative to medium baseline ± standard errors. The preferential impact of decline walking on braking forces and incline walking on propulsion ones was also observed in the after-effects. Namely, the flat group had smaller after-effects in braking than the decline group and in propulsion than the incline group.

**Figure 6.**
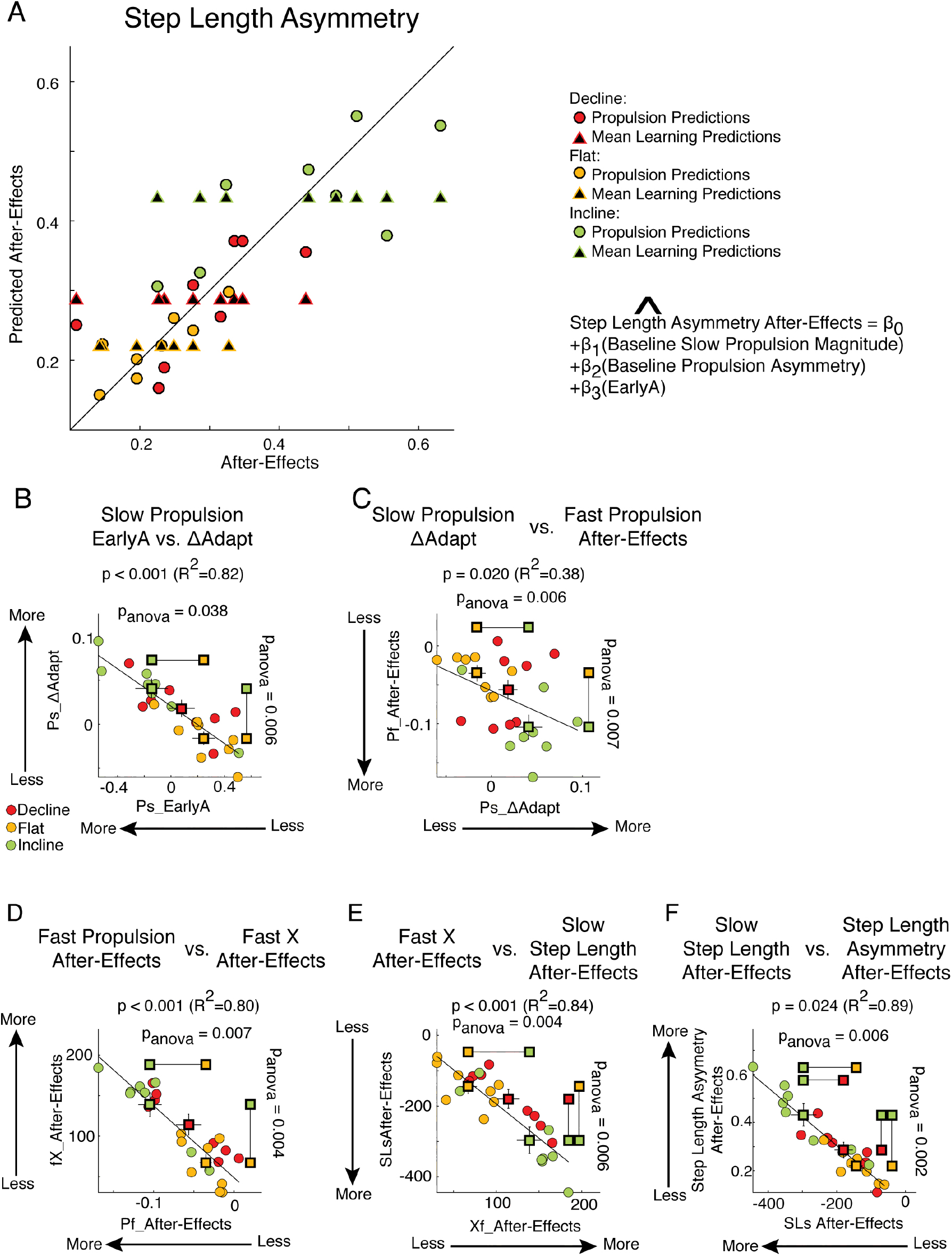
Propulsion-based statistical model and kinetic vs. kinematic regressions. (A) Individual subjects’ after-effects in step length asymmetry (x-axis) were estimated with two statistical models: the propulsion-based model and the mean-based model. The mean-based model estimated individual after-effects based on the sloped condition experienced by each subject. Conversely, the propulsion-based model estimated individual after-effects based on subject-specific propulsion during baseline and early adaptation according to the equation displayed at the top of the panel. The Propulsion-based model was able to account for significantly more inter-subject variance in after-effects of step length asymmetry than the mean-based model as indicated by a Bayes Factor analysis. (B-F) We show results from multiple or univariate linear regressions (performed when slope and continuous predictor were correlated). Values for individual subjects are shown with colored dots and group averages are illustrated with colored squares including standard errors. P-values from One-way ANOVAs comparing group means are listed parallel to each axis and significant differences between two groups according to post-hoc testing are indicated with lines connecting colored squares representing the groups that were different. (B) Slow leg’s propulsion during early adaptation vs. Ps_∆Adapt. Multiple regression indicated that the adaptation of the slow leg’s propulsion (Ps_∆Adapt) depended on the inclination condition and the slow leg’s propulsion during early adaptation. Specifically, large perturbations of the slow leg’s propulsion during early adaptation led to more adaptation of this leg’s propulsion force. (C) Slow leg’s propulsion adaptation (Ps_∆Adapt) vs. Fast leg’s propulsion post-adaptation after-effects (Pf_after-effects). Multiple regression indicated that the fast leg’s after-effects in propulsion depended on the inclination condition and the slow leg’s adaptation of propulsion forces. This contralateral relation between adaptation and post-adaptation was consistent with the one observed in step lengths of individual legs. (D) Positive correlation between after-effects in fast legs’ propulsion (Pf_after-effects) and fast leg’s X-position (fast X-position). In particular, subjects who generated smaller propulsion forces during post-adaptation were those whose trailing X-position was closer to their body (Large fast X-position values). This association is consistent with previous reports between magnitude of propulsion forces and trailing leg position (Leroux, et al 2002; Dewolf et al 2017). (E) Slow leg’s step length after-effects (SLs_after-effect) vs. Fast leg’s X-position. Multiple regression indicated that the after-effects in the slow leg’s step length depended on the inclination condition and the after-effects in the trailing leg’s position. More specifically, trailing X-positions close to the body (large X-positions) resulted in reduced step lengths (negative step length values) with respect to baseline walking. (F) Positive correlation between after-effects in step length asymmetry (SLA_after-effect) and step lengths of the leg that walked slow during the split condition (SLs_after-effect). This is consistent with results indicating that step length asymmetries during post-adaptation are mostly attributed to the slow leg’s step length after-effects.

Finally, we observed that this association between unilateral disturbance of slow leg’s propulsion and after-effects in step length asymmetry can be explained by the impact of propulsion forces on the slow step length during post-adaptation. This was indicated by regression analyses between kinetic and kinematic variables within and across adaptation and post-adaptation epochs (Figure 6B-F). These regressions revealed that large perturbations of the slow leg’s propulsion led to large adaptation of this parameter (Figure 6B; pmdl<0.001, R^2^=0.82), such that subjects that were perturbed the most were also those that adapted the slow leg’s propulsion the most. In addition, the adaptation of the slow leg’s propulsion force was positively associated with after-effects on the other leg’s propulsion (Figure 6C; p=0.020, R^2^=0.38). In other words, larger adaptation of the slow leg’s propulsion was related to greater after-effects on the propulsion of the fast side and not on the slow side (data not shown). This is consistent with the post-hoc observation that adaptation of step length on one side led to after-effects on the other side (see Figure 3). Importantly, there was a strong association between the fast leg’s propulsion after-effects and those of the trailing position of this leg (fast X-position) (Figure 6D; R^2^=0.80) when taking a step with the leg that walked slow in the split condition. Consistently, the after-effects of the fast leg’s X-position were associated to those of the slow leg’s step length (Figure 6E; p<0.001, R^2^=0.84), which was strongly associated with step length asymmetry after-effects (Figure 6F; R^2^=0.89). It is worth pointing out that group was a factor in all these regressions since it was either a significant categorical predictor in the multiple regressions (6B: Ps_EarlyA: p<0.001, pgroup<0.001; 6C: Ps_∆Adapt: p=0.68, pgroup=0.057; 6E: Xf_After-Effects: p<0.001, pgroup=0.015) or significantly correlated to the continuous variables (6D: Pearson p=0.043; 6F: Pearson p=0.024). This suggests that these associations might have been clearly observed in our study because of the range of inter-subject variability originated by the different sloped conditions. Taken together, the larger after-effects in step length asymmetry were mediated by unilateral step length after-effects on the side that walked slow, which was facilitated by large disturbance and adaptation of the slow leg’s propulsion.

## DISCUSSION

We investigated the influence of anterior-posterior forces on gait adaptation and learning induced by split-belt walking at different slopes, which naturally altered leg orientation and forces when feet were in contact with the ground. To our surprise, each inclination group recovered their baseline leg orientation at the expense of step length symmetry, which was a profound finding given that step length asymmetry is considered a biomarker of inefficient gait (Finley *et al.*, 2013; Bhounsule *et al.*, 2014; Awad *et al.*, 2015; Finley & Bastian, 2017). These distinct leg orientations were likely self-selected to generate the forces for walking at the specific speed and slope set by each split-belt task. This was achieved by distinct adaptation between the leading and trailing legs’ orientation at foot landing, suggesting the involvement of different physiological mechanisms in the control of leg orientation over the course of the stance phase. It was also unexpected that propulsion forces, rather than braking ones, influenced locomotor learning, which was indicated by the larger adaptation and after-effects of the sloped group with larger propulsion demands compared to the one with larger braking demands. The key role of propulsion forces was further supported by the fact that subject-specific propulsion tendencies during baseline and early adaptation were predictive of individual after-effects. Taken together, our findings suggest that altering propulsion forces during split-belt walking facilitates locomotor learning. Therefore, interventions augmenting propulsion demands could be more efficacious in the rehabilitation of hemiparetic gait.

### The motor system prioritizes the control of leg orientation over step length symmetry

All groups recovered their baseline leg orientation at the expense of step length symmetry, which indicated that speed-specific leg orientation was prioritized over symmetric step lengths. This has two key implications. First, step length symmetry is a gait feature that is not as valued by the motor system as previously considered. Specifically, step length symmetry is thought to be tightly controlled because subjects self-select symmetric step lengths even in asymmetric environments (e.g., Reisman *et al.*, 2005; Savin *et al.*, 2014) and those with more symmetric step lengths in these environments are also those spending less metabolic energy (Finley *et al.*, 2013). Thus, we surprisingly found that subjects in the incline and decline split-belt groups respectively overshot and undershot step length symmetry to recover baseline leg orientations. However, this finding can be explained by the impact of kinetic demands in shaping our movements, which is the second implication of our result. Notably, inverted pendulum models of walking suggest that subjects orient their legs to maintain a constant speed by equalizing positive and negative work over the gait cycle (Kajita *et al.*, 2001; Donelan *et al.*, 2002; Kuo, 2002; Kuo *et al.*, 2005). Consistently, we observed that subjects oriented their legs in the incline and decline groups to generate the forces counteracting the distinct effect of gravity on the center of mass at the different slopes (Lay *et al.*, 2006). Thus, leg orientation is closely regulated in order to walk at the distinct speeds and inclination (Leroux *et al.*, 2002; Orendurff *et al.*, 2008) imposed on each group. We particularly observed a strong effect of slope on leg orientation when walking slow, but this is also observed at fast speeds when using a coordinate frame aligned with gravity (Leroux *et al.*, 2002), rather than aligned with walking surface as done in this study. Given that humans have least-effort tendencies (Margaria, 1976; Alexander, 1989; Bertram & Ruina, 2001; Bertram, 2005; Kuo *et al.*, 2005; Selinger *et al.*, 2015) we think that step length symmetry is not necessarily energetically optimal in a split-belt environment (Sánchez *et al.*, 2017). Instead, subjects might self-select leg orientations to optimally generate the mechanical work (Selgrade *et al.*, 2017) for walking at the speed and inclination set on each foot, but further studies are needed to determine if this is the case.

The self-selected leg orientation also led to a contralateral relation between adaptation and post-adaptation. In other words, the disruption to forces and movements of individual legs led to more ipsilateral adaptation, but it did not result in larger ipsilateral after-effects. In fact, more adaptation of one leg changed the gait of the other leg as indicated by after-effects in kinetic and kinematic measures (e.g., propulsion forces, step length, and trailing position). This is in contrast to sensorimotor adaptation of other motor behaviors such as reaching, in which adaptation and de-adaptation effects are mostly constrained to a single effector (Nozaki *et al.*, 2006; Yokoi *et al.*, 2011, 2017; Wang *et al.*, 2013). The contralateral relation between adaptation and post-adaptation might be exclusive to locomotion; possibly because, unlike bimanual tasks (Ahmed *et al.*, 2008; Yokoi *et al.*, 2014), legs share a common goal in walking (i.e., keep the body on the treadmill) while experiencing opposite perturbations (i.e., one leg moves faster and the other one slower compared to regular walking) (Choi & Bastian, 2007). This contralateral relation between legs and unilateral effects reported on each group are, of course, occluded when using symmetry measures to characterize locomotor adaptation. Consequently, it is important to characterize the adaptation of each leg in tasks inducing locomotor learning (e.g., Reisman *et al.*, 2005), particularly when targeting unilateral deficits on hemiparetic gait (Bowden *et al.*, 2006; Balasubramanian *et al.*, 2007). Another consistent observation in the analysis of individual limbs was that all groups converged to the same step length value for the leg walking slow during adaptation. This is exclusively observed during split and not tied conditions, indicating that this might be task-constraint of split-belt walking, which will be the subject of future work. In sum, split-belt walking at different slopes predominantly altered the adaptation of one leg and let to subsequent after-effects on the other leg, highlighting the bilateral nature of sensorimotor adaptation in walking.

### Physiological mechanisms for controlling leg orientation

Our results suggest distinct control between the leading and trailing legs’ orientation when taking a step because they transition differently upon sudden changes in the walking environment. More specifically, the leading leg’s orientation at foot landing (α position) exhibits smooth and continuous changes when transitioning from the split to tied situations in all our inclination conditions and other perturbation magnitudes (Malone *et al.*, 2012), whereas the trailing leg’s orientation (X position) is discontinuous. This suggests that the leading leg’s orientation at foot landing is controlled in a feedforward manner–that is, it is planned before the movement is executed based on an a slowly updated internal representation of the environment. This is supported by the fact that sensory information about leg orientation at foot landing is sent to cerebellar structures (Bosco & Poppele, 2001), housing the feedforward control of movements (Herzfeld *et al.*, 2015) and the fact that spinalized cats cannot adapt the orientation of the leading leg during split-belt walking (Frigon *et al.*, 2017). On the other hand, the trailing leg’s orientation when taking a step could be determined by a combination of feedback and feedforward control. Consider that feedback mechanisms adjust our movements by transforming delayed sensory information into actions in real-time (Jordan & Rumelhart, 1992; Bhushan & Shadmehr, 1999). Accordingly, the trailing leg’s orientation is immediately regulated upon manipulations to ipsilateral sensory information from spindles in hip muscles, load sensors and cutaneous information (Grillner & Rossignol, 1978; Duysens & Pearson, 1980; Duysens *et al.*, 2000; Pang & Yang, 2000; Rossignol, 2006). In addition, the trailing leg’s orientation is determined by the step time (i.e., period between ipsilateral and contralateral heel-strikes) (Finley *et al.*, 2015), which is controlled in a feedforward manner as suggested by behavioral split-belt studies (Malone *et al.*, 2012; Finley *et al.*, 2015) and Purkinjie cells tracking heel-strikes (Apps *et al.*, 1995; Bosco & Poppele, 2001). In particular, the feedforward control of step timing and hence that of the trailing leg’s orientation might be based on subjects’ expectation of the speed at which the belt is moving. Taken together, our results suggest distinct control mechanisms of the leading and trailing legs’ orientation when taking a step: the leading position is mostly mediated by feedforward mechanisms, whereas the trailing position is mediated by a combination of feedforward and feedback mechanisms.

### Motor adaptation and learning is facilitated in walking when manipulating propulsion forces, rather than braking ones

While braking and propulsion forces were adapted in all groups, augmenting propulsion forces facilitated motor adaptation. This was indicated by the greater adaptation and after-effects of the incline group with larger propulsion demands, whereas the adaptation was reduced and the after-effects were unchanged in the decline group with larger braking demands. The preferential impact of propulsion on locomotor adaptation was unexpected given prior studies reporting minimal (Roemmich *et al.*, 2012) or absent adaptation of propulsion forces (Ogawa *et al.*, 2014) and subsequently lacking propulsion after-effects (Roemmich *et al.*, 2012; Ogawa *et al.*, 2014) in contrast with the adaptation and after-effects of braking forces (Ogawa *et al.*, 2014). Our findings are at odds with these reports possibly because our participants experienced greater speed differences and were adapted at a more naturalistic walking speed (i.e., mean speed across legs of 1m/s vs. 0.75m/s). We additionally found that the incline group adapted faster, indicating that inclined walking also augmented the saliency (i.e., easier to detect) and/or sensitivity (i.e., quicker to respond) to the split-belt perturbation. The saliency of the speed difference between legs could be augmented in the inclined condition because its reduced cadence (Kawamura *et al.*, 1991; Sun *et al.*, 1996; McIntosh *et al.*, 2006; Phan *et al.*, 2013)(and hence longer stance times (i.e., period in direct contact with the environment), which increase subjects’ ability to perceive speed differences (Hoogkamer *et al.*, 2015). In addition, sensory inputs encoding walking speed are stimulated more when walking incline (Sinkjaer *et al.*, 2000; Lamont & Zehr, 2006; Klint *et al.*, 2008; Tillakaratne *et al.*, 2014; Choi *et al.*, 2016) also increasing the saliency of speed differences between the legs in the split situation. On the other hand, subjects in the split incline group might have adapted faster because they were more sensitive to increments in energetic cost due to step length asymmetry (Finley *et al.*, 2013; Bhounsule *et al.*, 2014), because incline walking has large energetic demands in and of itself (Johnson *et al.*, 2002). Therefore, larger propulsion demands experienced by the incline group resulted in more and faster gait adaptation, indicating that altering propulsion forces during split-belt walking facilitated locomotor learning.

The relevance of propulsion forces was also evident by the association between individual tendencies in propulsion forces during baseline and early adaptation and subject-specific after-effects. The positive relation between large movement disruptions in early adaptation and after-effects is well documented in error-based learning (Körding & Wolpert, 2004; Wei & Körding, 2009; Green *et al.*, 2010). Conversely, the predictive nature of baseline features and motor learning is less commonly observed in sensorimotor adaptation (Kelly & Sober, 2014; Wu *et al.*, 2014). Here we find that individuals biased to propel more or asymmetrically during baseline walking exhibit larger after-effects. We speculate that these baseline features possibly led to more adaptation and consequently larger after-effects. Consider that the split perturbation disrupts propulsion away from the baseline behavior and subjects adapt towards their original bias, as observed in other gait features (Malone & Bastian, 2014). Therefore, individuals with greater biases need to adapt more to approach their larger baseline bias. In addition, we observed that the asymmetric bias was in the same direction as the one observed during late adaptation. Thus, subjects that were more asymmetric might adapt more because they are more prone to adopt the asymmetric pattern needed to walk in the split condition. Of note, we observed a large range of propulsion forces during baseline and early adaptation across individuals, which might be needed to identify the reported association between propulsion forces and after-effects. In sum, our findings indicate that augmenting propulsion demands during split-belt walking facilitates gait adaptation and locomotor learning, suggesting that altering propulsion forces could be used as a training stimulus for gait rehabilitation.

#### Clinical implications

There is an interest in modulating the extent of split-belt adaptation and learning as stroke subjects tend to only adapt to their baseline asymmetry during adaptation (Malone & Bastian, 2014) and do not always respond to split-belt training interventions (Reisman *et al.*, 2013). Our findings suggest that tasks disrupting propulsion could be used as a strategy for facilitating motor learning to correct pathological gait. For example, incline spit-belt walking could lead to more adaptation and after-effects in post-stroke individuals (Sombric *et al.*, 2015). Importantly, incline split-belt walking requires more propulsion of the side walking slow in the asymmetric environment. Thus, incline split-belt walking could reinforce two target behaviors of the paretic side if placed in the slow belt: 1) longer stance time duration and 2) greater propulsion forces, both of which result in post-stroke gait asymmetry (Bowden *et al.*, 2006; Balasubramanian *et al.*, 2007). Moreover, we find that incline split-belt walking leads to faster adaptation. This could be exploited to increase the adaptation rate in older populations (Sombric *et al.*, 2017), which could benefit the training of older clinical populations. Our results also indicate that baseline features of individual subjects are informative about subject-specific capacity to modify their gait and learn new walking patterns, which is critical for understanding inter-subject differences in motor corrections post-stroke. Future work will be needed to determine the efficacy of incline split-belt walking over other split-belt training protocols (Reisman *et al.*, 2013; Lewek *et al.*, 2017), including the assessment of whether corrected behavior in the incline condition carries over to walking in a level surface over ground.

## Author Contributions

All experiments were performed in the Sensorimotor Learning Laboratory. G.T. and C.S. were involved with the conception and design of the work. C.S. and J.C. collected and analyzed the data. C.S and G.T. interpreted the results. C. S. drafted the manuscript, which was carefully revised by all authors. The final version of the manuscript has been approved by all the authors who agree to be accountable for all aspects of the work in ensuring that questions related to the accuracy or integrity of any part of the work are appropriately investigated and resolved. All authors qualify for authorship and all those who qualify for authorship are listed.

## Funding

C.S. is funded by a fellowship from the National Science Foundation (NSF-GRFP).

CRDF 4.30204; NSF1535036; AHA 15SDG25710041

## Acknowledgements

The authors acknowledge the valuable input from Pablo Iturralde and Digna de Kam.

## Bibliography

Ahmed AA, Wolpert DM & Flanagan JR (2008). Flexible Representations of Dynamics Are Used in Object Manipulation. Curr Biol 18, 763–768.

Alexander RM (1989). Optimization and gaits in the locomotion of vertebrates. Physiol Rev 69, 1199–1227.

Apps R, Hartell NA & Armstrong DM (1995). Step phase-related excitability changes in spino-olivocerebellar paths to the c1 and C3 zones in cat cerebellum. 687–702.

Awad LN, Palmer JA, Pohlig RT, Binder-Macleod SA & Reisman DS (2015). Walking speed and step length asymmetry modify the energy cost of walking after stroke. Neurorehabil Neural Repair 29, 416–423.

Balasubramanian CK, Bowden MG, Neptune RR & Kautz SA (2007). Relationship Between Step Length Asymmetry and Walking Performance in Subjects With Chronic Hemiparesis. Arch Phys Med Rehabil 88, 43–49.

Bertram JEA (2005). Constrained optimization in human walking: cost minimization and gait plasticity. J Exp Biol 208, 979–991.

Bertram JEA & Ruina A (2001). Multiple walking speed-frequency relations are predicted by constrained optimization. J Theor Biol 209, 445–453.

Beschorner KE, Albert DL & Redfern MS (2016). Required coefficient of friction during level walking is predictive of slipping. Gait Posture 48, 256–260.

Bhounsule PA, Cortell J, Grewal A, Hendriksen B, Karssen JGD, Paul C & Ruina A (2014). Low-bandwidth reflex-based control for lower power walking : 65 km on a single.; DOI: 10.1177/0278364914527485.

Bhushan N & Shadmehr R (1999). Evidence for a forward dynamics model in human adaptive motor control. Adv Neural Inf Process Syst 11, 3–9.

Bosco G & Poppele RE (2001). Proprioception from a spinocerebellar perspective. Physiol Rev 81, 539–568.

Bowden MG, Balasubramanian CK, Neptune RR & Kautz S a. (2006). Anterior-Posterior Ground Reaction Forces as a Measure of Paretic Leg Contribution in Hemiparetic Walking. Stroke 37, 872–876.

Bruijn SM, Van Impe A, Duysens J & Swinnen SP (2012). Split-belt walking: adaptation differences between young and older adults. J Neurophysiol 108, 1149–1157.

Choi JT & Bastian AJ (2007). Adaptation reveals independent control networks for human walking. Nat Neurosci 10, 1055–1062.

Choi JT, Jensen P, Nielsen JB & Bouyer LJ (2016). Error signals driving locomotor adaptation : cutaneous feedback from the foot is used to adapt movement during perturbed walking. J Physiol 19, 5673–5684.

Dewolf AH, Ivanenko YP, Lacquaniti F & Willems PA (2017). Pendular energy transduction within the step during human walking on slopes at different speeds. 1–25.

Donelan JM, Kram R & Kuo AD (2002). Simultaneous positive and negative external mechanical work in human walking. J Biomech 35, 117–124.

Duysens J, Clarac F, Cruse H, Fysica M, National C & Recherche D (2000). Load-regulating mechanisms in gait and posture: comparative aspects. Physiol Rev 80, 83–133.

Duysens J & Pearson K (1980). Inhibition of flexor burst generation by loading ankle extensor muscles in walking cats. Brain Res 187, 321–332.

Finley JM & Bastian AJ (2017). Associations Between Foot Placement Asymmetries and Metabolic Cost of Transport in Hemiparetic Gait.; DOI: 10.1177/1545968316675428.

Finley JM, Bastian AJ & Gottschall JS (2013). Learning to be economical: the energy cost of walking tracks motor adaptation. J Physiol 591, 1081–1095.

Finley JM, Long A, Bastian AJ & Torres-Oviedo G (2015). Spatial and Temporal Control Contribute to Step Length Asymmetry During Split-Belt Adaptation and Hemiparetic Gait. Neurorehabil Neural Repair 29, 786–795.

Frigon A, Thibaudier Y, Hurteau M & Dambreville C (2017). Left – right coordination from simple to extreme conditions during split-belt locomotion in the chronic spinal adult cat. 1, 341–361.

Green DA, Bunday KL, Bowen J, Carter T & Bronstein AM (2010). What does autonomic arousal tell us about locomotor learning? 170, 42–53.

Grillner S & Rossignol S (1978). On the initiation of the swing phase of locomotion in chronic spinal cats. Brain Res 146, 269–277.

Herzfeld DJ, Kojima Y, Soetedjo R & Shadmehr R (2015). Encoding of action by the Purkinje cells of the cerebellum. Nature 526, 439–441.

Hoogkamer W, Bruijn SM, Potocanac Z, Van Calenbergh F, Swinnen SP & Duysens J (2015). Gait asymmetry during early split-belt walking is related to perception of belt speed difference. J Neurophysiol 114, 1705–1712.

Item-glatthorn JF, Casartelli NC & Maffiuletti NA (2016). Reproducibility of gait parameters at different surface inclinations and speeds using an instrumented treadmill system. 44, 259–264.

Johnson AT, Benhur Benjamin M & Silverman N (2002). Oxygen consumption, heat production, and muscular efficiency during uphill and downhill walking. Appl Ergon 33, 485–491.

Jordan MI & Rumelhart DE (1992). Forward models{\textasciitilde}: supervised learning with a distal teacher. Cogn Sci 307354, 16:307––354.

Jørgensen CK, Fink P & Olesen F (2000). Psychological distress among patients with musculoskeletal illness in general practice. Psychosomatics 41, 321–329.

Kajita S, Kanehiro F, Kaneko K, Yokoi K & Hirukawa H (2001). The 3D linear inverted pendulum mode: a simple modeling for a biped walking pattern generation. Proc 2001 IEEE/RSJ Int Conf Intell Robot Syst Expand Soc Role Robot Next Millenn (Cat No01CH37180) 1, 239–246.

Kass RE, Raftery AE, Kass RE & Raftery AE (1995). Bayes Factors Bayes Factors. 90, 773–795.

Kawamura K, Tokuhiro A & Takechi H (1991). Gait analysis of slope walking: a study on step length, stride width, time factors and deviation in the center of pressure. Acta Med Okayama 45, 179–184.

Kelly CW & Sober SJ (2014). A simple computational principle predicts vocal adaptation dynamics across age and error size. Front Integr Neurosci 8, 1–9.

Klint RA, Nielsen JB, Cole J, Sinkjaer T & Grey MJ (2008). Within-step modulation of leg muscle activity by afferent feedback in human walking. J Physiol 586, 4643–4648.

Körding KP & Wolpert DM (2004). The loss function of sensorimotor learning. Proc Natl Acad Sci U S A 101, 9839–9842.

Kuo AD (2002). Energetics of Actively Powered Locomotion Using the Simplest Walking Model. J Biomech Eng 124, 113.

Kuo AD, Donelan JM & Ruina A (2005). Energetic Consequences of Walking Like an Inverted Pendulum : Step-to-Step Transitions.

Lamont E V. & Zehr EP (2006). Task-specific modulation of cutaneous reflexes expressed at functionally relevant gait cycle phases during level and incline walking and stair climbing. Exp Brain Res 173, 185–192.

Lay AN, Hass CJ & Gregor RJ (2006). The effects of sloped surfaces on locomotion : A kinematic and kinetic analysis. 39, 1621–1628.

Lay AN, Hass CJ, Nichols TR & Gregor RJ (2007). The effects of sloped surfaces on locomotion : An electromyographic analysis. 40, 1276–1285.

Leroux A, Fung J & Barbeau H (2002). Postural adaptation to walking on inclined surfaces : I. Normal strategies. Gait Posture 15, 64–74.

Lewek MD, Bradley CE, Wutzke CJ & Zinder^ SM (2014). The Relationship Between Spatiotemporai Gait Asymmetry and Balance in Individuals With Chronic Stroke. J Appl Biomech 30, 31–36.

Lewek MD, Braun CH & Wutzke C (2017). The role of movement errors in modifying spatiotemporal gait asymmetry post stroke : a randomized controlled trial.; DOI: 10.1177/0269215517723056.

Malone L, Bastian A & Torres-Oviedo G (2012). How does the motor system correct for errors in time and space during locomotor adaptation? J Neurophysiol 108, 672–683.

Malone LA & Bastian AJ (2014). Spatial and Temporal Asymmetries in Gait Predict Split-Belt Adaptation Behavior in Stroke.; DOI: 10.1177/1545968313505912.

Margaria R (1976). Biomechanics and Energetics of Muscular Exercise. Clarendon Press, Oxford, UK.

Mawase F, Haizler T, Bar-Haim S & Karniel A (2013). Kinetic adaptation during locomotion on a split-belt treadmill. J Neurophysiol 109, 2216–2227.

McIntosh AS, Beatty KT, Dwan LN, Vickers DR, Ã ASM, Beatty KT, Dwan LN & Vickers DR (2006). Gait dynamics on an inclined walkway. J Biomech 39, 2491–2502.

Nozaki D, Kurtzer I & Scott SH (2006). Limited transfer of learning between unimanual and bimanual skills within the same limb. Nat Neurosci 9, 1364–1366.

Ogawa T, Kawashima N, Ogata T & Nakazawa K (2014). Predictive control of ankle stiffness at heel contact is a key element of locomotor adaptation during split-belt treadmill walking in humans. J Neurophysiol 111, 722–732.

Orendurff MS, Bernatz GC, Schoen JA & Klute GK (2008). Kinetic mechanisms to alter walking speed. Gait Posture 27, 603–610.

Pang MYC & Yang JF (2000). The initiation of the swing phase in human infant stepping: Importance of hip position and leg loading. J Physiol 528, 389–404.

Patterson KK, Nadkarni NK, Black SE & McIlroy WE (2012). Gait symmetry and velocity differ in their relationship to age. Gait Posture 35, 590–594.

Phan PL, Blennerhassett JM, Lythgo N, Dite W & Morris ME (2013). Over-ground walking on level and sloped surfaces in people with stroke compared to healthy matched adults. Disabil Rehabil 35, 1302–1307.

Redfern MS, Cham R, Gielo-Perczak K, Grönqvist R, Hirvonen M, Lanshammar H, Marpet M, Pai IV CY-C & Powers. C (2001). Biomechanics of slips. Biomechanics 44, 1138–1166.

Reisman DS, Block HJ, Bastian AJ, Darcy S, Block HJ & Inter- AJB (2005). Interlimb coordination during locomotion: what can be adapted and stored? J Neurophysiol 94, 2403–2415.

Reisman DS, McLean H, Keller J, Danks KA & Bastian AJ (2013). Repeated split-belt treadmill training improves poststroke step length asymmetry. Neurorehabil Neural Repair 27, 460–468.

Reisman DS, Wityk R, Silver K & Bastian AJ (2009). Split-belt treadmill adaptation transfers to overground walking in persons poststroke. Neurorehabil Neural Repair 23, 735–744.

Roemmich RT, Stegemöller EL & Hass CJ (2012). Lower extremity sagittal joint moment production during split-belt treadmill walking. J Biomech 45, 2817–2821.

Rossignol S (2006). Dynamic Sensorimotor Interactions in Locomotion. Physiol Rev 86, 89–154.

Sánchez N, Park S & Finley JM (2017). Evidence of Energetic Optimization during Adaptation Differs for Metabolic, Mechanical, and Perceptual Estimates of Energetic Cost. 1–14.

Savin DN, Morton SM & Whitall J (2014). Generalization of improved step length symmetry from treadmill to overground walking in persons with stroke and hemiparesis. Clin Neurophysiol 125, 1012–1020.

Selgrade BP, Thajchayapong M, Lee GE, Toney ME & Chang Y-H (2017). Changes in mechanical work during neural adaptation to asymmetric locomotion. J Exp Biol 220, 2993–3000.

Selinger JC, Shawn M, Connor O, Wong JD, Maxwell J, Selinger JC, Connor SMO, Wong JD & Donelan JM (2015). Humans Can Continuously Optimize Energetic Cost during Walking Report Humans Can Continuously Optimize Energetic Cost during Walking. Curr Biol 25, 2452–2456.

Sinkjaer T, Andersen JB, Ladouceur M, Christensen LO & Nielsen JB (2000). Major role for sensory feedback in soleus EMG activity in the stance phase of walking in man. J Physiol 523 Pt 3, 817–827.

Sombric CJ, Harker HM, Sparto PJ & Torres-oviedo G (2017). Explicit Action Switching Interferes with the Context-Specificity of Motor Memories in Older Adults. Front Aging Neurosci; DOI: 10.3389/fnagi.2017.00040.

Sombric CJ, Mariscal DM, Calvert JS, Iturralde PA & Torres-Oviedo G (2015). It’s all uphill from here: incline split-belt walking increases locomotor learning post-stroke. In Advances in Motor Learning & Motor Control.

Sun J, M. W, Svensson N & Lloyd, D. (1996). 1996. The influence of surface slope on human gait characteristics: a study of urban pedestrians walking on an inclined surface. Ergonomics 39, 677–692.

Tillakaratne NJK, Duru P, Fujino H, Zhong H, Xiao MS, Edgerton VR & Roy RR (2014). Identification of Interneurons Activated at Different Inclines During Treadmill Locomotion in Adult Rats. J Neuroscince Res 92, 1714–1722.

Torres-Oviedo G & Bastian AJ (2010). Seeing Is Believing : Effects of Visual Contextual Cues on Learning and Transfer of Locomotor Adaptation. J Neurosci 30, 17015–17022.

Torres-Oviedo G & Bastian AJ (2012). Natural error patterns enable transfer of motor learning to novel contexts. J Neurophysiol 107, 346–356.

Wagenmakers EJ (2007). A practical solution to the pervasive problems of p values. Psychon Bull Rev 14, 779–804.

Wang J, Lei Y, Xiong K & Marek K (2013). Substantial Generalization of Sensorimotor Learning from Bilateral to Unilateral Movement Conditions. PLoS One; DOI: 10.1371/journal.pone.0058495.

Waters RL & Mulroy S (1999). The energy expenditure of normal and pathologic gait (ABS). Gait Posture 9, 207–231.

Wei K & Körding K (2009). Relevance of error: what drives motor adaptation? J Neurophysiol 101, 655–664.

Wu HG, Miyamoto YR, Nicolas L, Castro G, Smith MA & Biology E (2014). Temporal structure of motor variability is dynamically regulated and predicts motor learning ability Howard. Nat Neurosci 17, 312–321.

Yokoi A, Bai W & Diedrichsen J (2017). Restricted transfer of learning between unimanual and bimanual finger sequences. J Neurophysiol 117, 1043–1051.

Yokoi A, Hirashima M & Nozaki D (2011). Gain Field Encoding of the Kinematics of Both Arms in the Internal Model Enables Flexible Bimanual Action. J Neurosci 31, 17058–17068.

Yokoi A, Hirashima M & Nozaki D (2014). Lateralized Sensitivity of Motor Memories to the Kinematics of the Opposite Arm Reveals Functional Specialization during Bimanual Actions. J Neurosci 34, 9141–9151.

Yokoyama H, Sato K, Ogawa T, Yamamoto S-I, Nakazawa K & Kawashima N (2018). Characteristics of the gait adaptation process due to split-belt treadmill walking under a wide range of right-left speed ratios in humans. PLoS One 13, e0194875.

